# Functional characterization and discovery of modulators of SbMATE, the agronomically important aluminium tolerance transporter from *Sorghum bicolor*

**DOI:** 10.1101/182964

**Authors:** Rupak Doshi, Aaron P. McGrath, Miguel Piñeros, Paul Szewczyk, Denisse M. Garza, Leon V. Kochian, Geoffrey Chang

**Affiliations:** Skaggs School of Pharmacy and Pharmaceutical Sciences, University of California at San Diego, La Jolla, California, USA.; Robert W. Holley Center for Agriculture and Health, United States Department of Agriculture–Agricultural Research Service, Cornell University, Ithaca, NY, USA; Global Institute for Food Security, University of Saskatchewan, Saskatoon, Canada; Department of Pharmacology, School of Medicine, University of California at San Diego, La Jolla, California, USA

## Abstract

About 50% of the world’s arable land is strongly acidic (soil pH < 5). The low pH of these soils solubilizes root-toxic ionic aluminium (Al^3+^) species from clay minerals, driving the evolution of various counteractive adaptations in cultivated crops. The food crop *Sorghum bicolor*, for example, upregulates the membrane-embedded transporter protein SbMATE in its roots. SbMATE mediates efflux of the anionic form of the organic acid, citrate, into the soil rhizosphere, chelating Al^3+^ ions and thereby imparting Al-resistance based on excluding Al^+3^ from the growing root tip. Here, we use electrophysiological, radiolabeled, and fluorescence-based transport assays in two heterologous expression systems to establish a broad substrate recognition profile of SbMATE, showing the transport of ^14C^- citrate anion, as well as the organic monovalent cation, ethidium, but not the divalent ethidium-derivative, propidium. The transport cycle is proton and/or sodium-driven, and shares certain molecular mechanisms with bacterial MATE-family transporters. We further complement our transport assays by directly measuring substrate binding to detergent-purified SbMATE protein. Finally, we use the functionally-folded, purified membrane protein as an antigen to discover high-affinity, native conformation-binding and transport function-altering nanobodies using an animal-free, mRNA/cDNA display technology. Our results demonstrate the utility of using *Pichia pastoris* as an efficient eukaryotic host to express large quantities of functional plant transporter proteins for *in vitro* characterization. The nanobody discovery approach is applicable to other low immunogenic plant proteins.

## INTRODUCTION

Nearly 30% of all cellular genomes code for integral membrane proteins, such as receptors, channels, and transporters. These proteins contain multiple membrane-spanning segments that traverse the phospholipid bilayer. Transporters are cellular gate-keepers, moving a wide variety of specific molecules and ions in both directions across the plasma membrane as well as other endomembrane-bound compartments. Importers allow the cellular uptake of essential nutrients and metabolic reactants, while exporters mediate the efflux of metabolic byproducts, defense agents, toxic compounds, and xenobiotics. As a result, transporters are crucial in shaping the ability of cells to adapt and survive in their surrounding environment.

Active transporters move substrates against their concentration or electrochemical (for ions) gradients utilizing energy, derived either from the binding and hydrolysis of ATP (primary active) or from the transmembrane electrochemical potential gradient (secondary active). The <u>M</u>ultidrug <u>A</u>nd <u>T</u>oxic compound <u>E</u>xtrusion (or MATE) family of secondary-active transporters was initially identified from bacterial isolates that appeared to be resistant to antibiotics, drugs, and cationic dyes. Since then, MATE transporters have been identified in almost all organisms, including humans and plants. In humans, MATE transporters play a key role in secreting organic drugs, toxins, and metabolites into the renal tubular lumen (*1*). In plants, MATE transporters are involved in a wide variety of physiological process, including xenobiotic efflux, accumulation of secondary metabolites including alkaloids and flavonoids, iron (Fe) translocation, plant hormone signaling, and aluminum (Al^3+^) resistance (*2*).

Given that aluminum (Al) is the most abundant metal in the earth’s crust, plants have evolved in a soil environment where their root system can frequently be exposed to high levels of this toxic metal. Acid soils constitute approximately 30% of the land presently under cultivation and over 50% of the world’s potentially arable lands (*3–4*). At soil pH <5.0, Al solubilizes as the phytotoxic species, Al^3+^, causing a strong inhibition of root growth and function, ultimately resulting in severe reductions in crop yield. Aluminum toxicity is, therefore, a significant global agricultural problem, prevalent in many regions where food security is already tenuous, with 38% of the farmland in Southeast Asia, 31% of Latin America, and 20% of the arable lands in East Asia and sub-Saharan Africa currently being impacted. The ability to adapt/fortify crops to tolerate high soil Al thus has the potential to enormously increase food production in many at-risk regions.

*Sorghum bicolor* is the fifth most widely grown cereal crop in the world, with the largest producers being Nigeria, India, USA, and Mexico. By studying genotypic differences between sorghum species that are sensitive or resistant to aluminum, a MATE family transporter, SbMATE, was identified via the positional cloning of the major sorghum Al resistance locus (*5*). SbMATE is highly upregulated in Al-tolerant sorghum, and in response to increased Al^3+^ levels in the soil, it underlies an increase in organic anion citrate exudation from roots into the surrounding rhizosphere, thereby chelating the free Al^3+^, forming nontoxic complexes that are no longer taken up by the roots. Consistently, transgenic SbMATE was also found to confer Al-tolerance in barley, which is one of the most Al-sensitive cereal crop species (*6*).

Despite belonging to the MATE family of proteins that are known to transport multiple chemically-diverse compounds, SbMATE and other Al-resistance associated orthologues (*5, 7–13*) have been studied purely from the perspective of their organic acid anion (i.e. citrate) efflux activity (Fig. 1A). Thus, a significant gap exists in our understanding of other classes of substrates that could potentially be transported. Furthermore, investigating the molecular mechanisms that govern SbMATE mediated transport is critical towards devising strategies that could improve aluminum tolerance traits in crops expressing SbMATE-type transporters in their roots.

**Figure 1:**
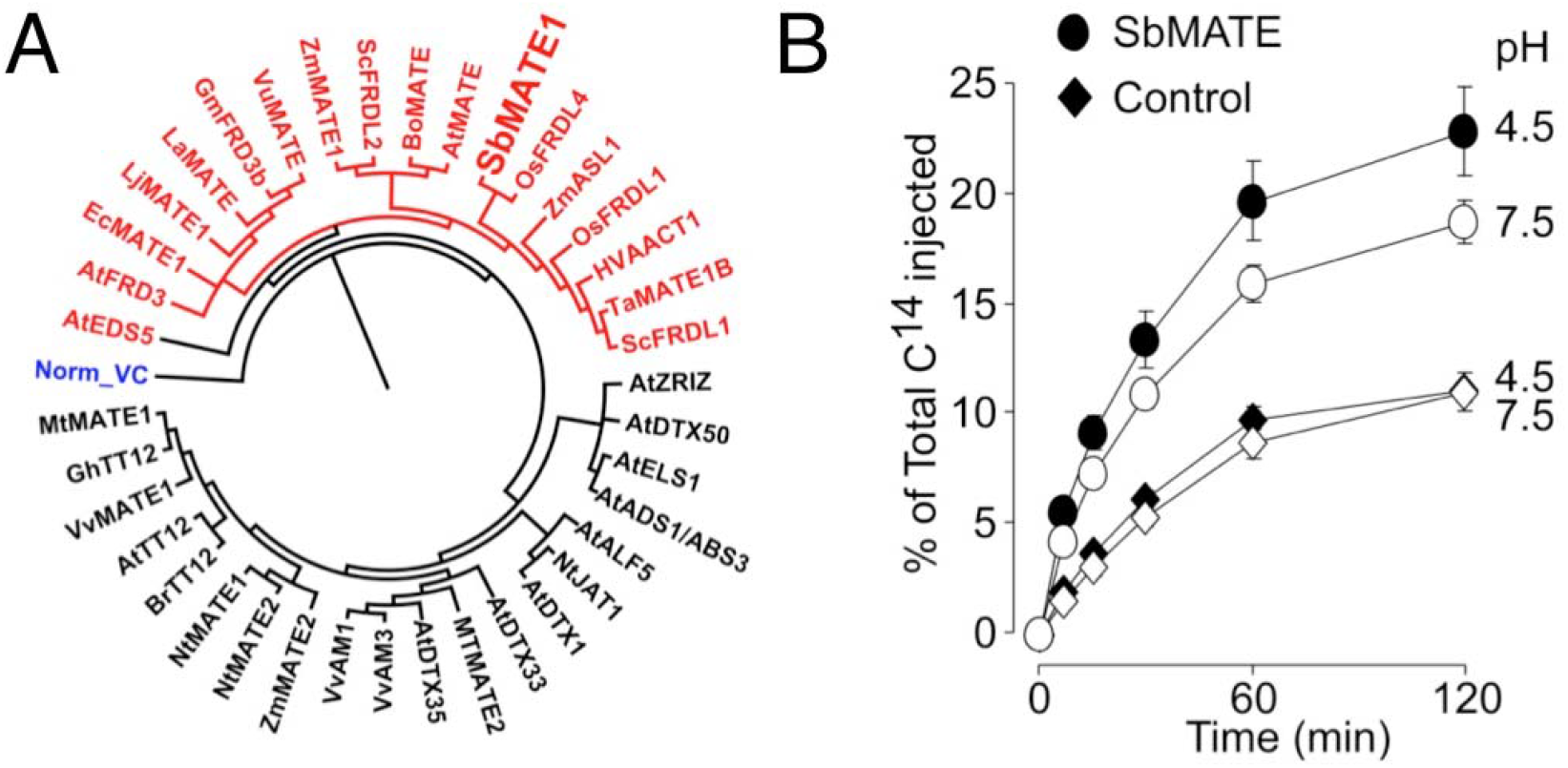
Phylogenetic analysis of plant MATE transporters and biophysical characterization of SbMATE-mediated citrate transport. (A) Phylogenetic analysis of the functionally characterized plant MATE transporters. Plant MATEs colored in red represent MATEs associated with Al-resistance and Fe transport/homeostasis. The bacterial NorM from *Vibrio cholerae* is shown for reference and it is colored in blue. The tree was built using protein sequences in Geneious Tree Builder. (B) SbMATE mediates citrate efflux in oocytes. Control (diamond symbols) and SbMATE (circle symbols) expressing cells were microinjected with [^14^C] citrate and kept in Ringer solution at the pH values indicated on the right margin (pH 7.5 and 4.5; open and filled symbols, respectively). The radioactivity in the bathing solutions was measured at the indicated time points; values are expressed as a percentage of the total radioactivity injected.

Here, we have attained a broader understanding of SbMATE’s functionality by studying its transport characteristics in a series of biochemical and biophysical assays, using cell-based transport assays in two different heterologous expression systems, and also in detergent-purified liposomal-reconstituted formulations. Additionally, we have used purified SbMATE as a model, plant-derived, full-length integral membrane protein to discover single-domain antibody reagent tools that bind the native, folded conformation of SbMATE and affect its transport characteristics.

## RESULTS

### SbMATE exhibits broad substrate recognition – organic anion (citrate) and cation (ethidium) transport

We have previously reported the use of Xenopus oocytes to express SbMATE and measure its electrogenic transport activity (*5, 8*). We used this heterologous expression system to directly measure the efflux of ^14^C-citrate from pre-loaded cells in the presence of an inwardly-directed proton gradient, i.e. substrate-proton antiport. Consistent with previous *in planta* experiments using transgenic *A. thaliana* lines, SbMATE-expressing oocytes mediated ^14^C-citrate efflux, quantified as ~2.5-fold higher ^14^C-citrate released from loaded cells relative to the basal levels from non-expressing control cells (Fig. 1B).

We further investigated substrate recognition and the proton antiport mechanism of SbMATE in *Pichia pastoris*. Intact yeast cells expressing SbMATE, empty vector-transformed control cells, and their respective uninduced control cells were washed and maintained on ice in phosphate-saline (PBS) buffer, pH 7.4. For the transport measurements, cells were incubated with 50 µM of ethidium bromide in buffer salt-concentration matched to PBS at pH 8.5 (i.e. 137 mM NaCl, 2.7 mM KCl). In this reversed, outwardly-directed proton gradient (assuming a near neutrality cytosolic pH; (*14*)), SbMATE exhibited uptake of ethidium into cells, as shown by a large increase in fluorescence (caused by the binding of ethidium to DNA) in the induced SbMATE-expressing cells, compared to control nonexpressing, and uninduced cells (Fig. 2A, 2B). We further measured the substrate-concentration dependence of SbMATE-mediated ethidium transport, which resulted in a halfmaximal transport rate computed at 130 ±17.8 µM ethidium bromide (mean ± SEM, n=3; Fig. 2C).

**Figure 2:**
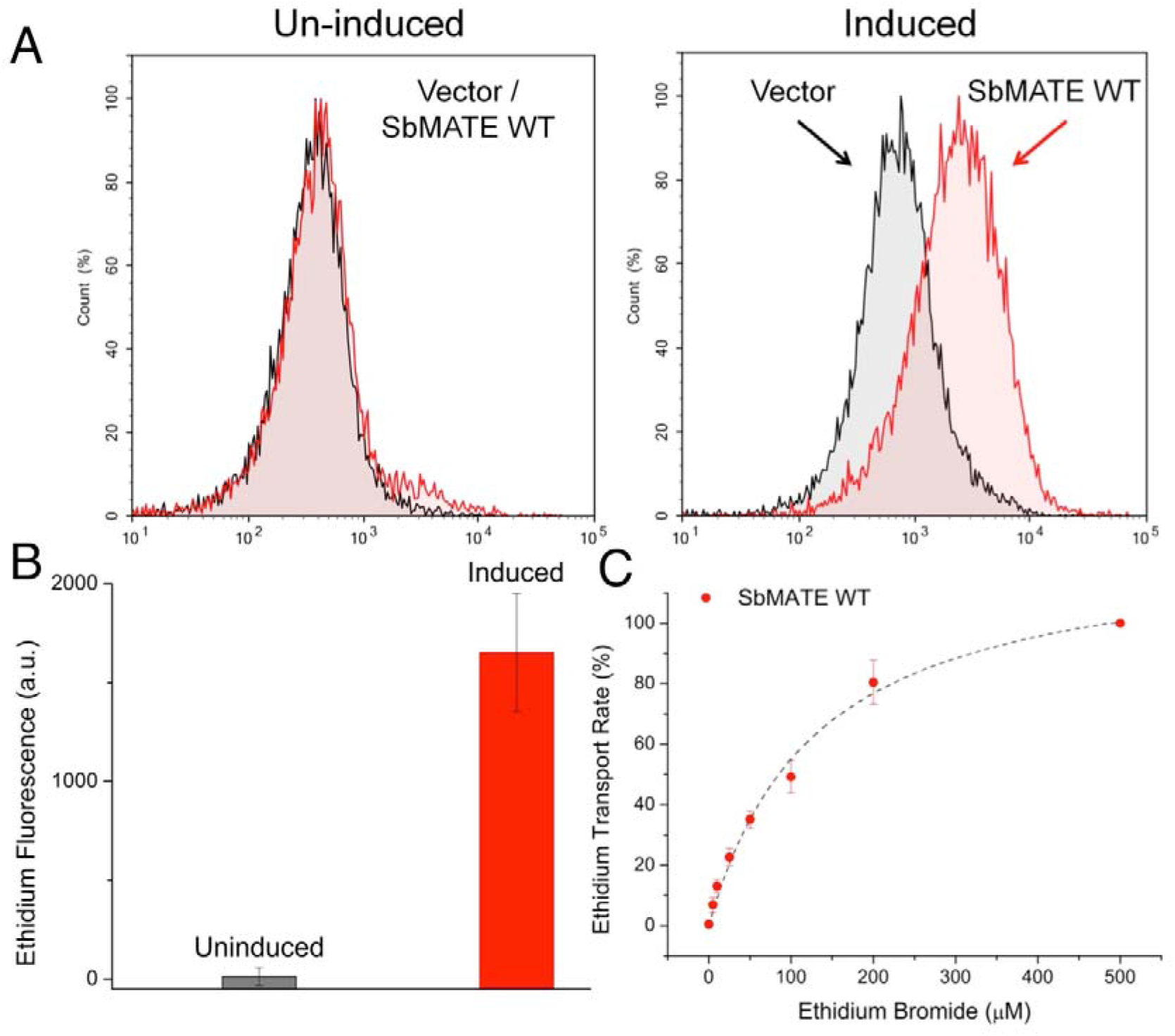
SbMATE expression in *P. pastoris* and transport of the monovalent organic cation, ethidium. (A) Uninduced and induced cells transformed with SbMATE WT or vector control were normalized to an OD_600_ of 15, washed, and maintained in standard PBS buffer, pH 7.4. For the ethidium transport assay, 50 μM ethidium bromide and a pH 8.5 buffer (25 mM Tris-Cl, pH 8.5, 137 mM NaCl, 2.7 mM KCl) was used to generate an outward-proton gradient. Ethidium uptake was measured over a 20 min uptake period using flow cytometry, with 12,000 events collected per sample. Histograms were plotted for ethidium fluorescence vs normalized frequency (n=5 replicates conducted with independently grown batches of cells, histograms shown are from a single experiment). (B) Median fluorescence intensities of uninduced or induced cells transformed with SbMATE WT were subtracted from the respective vector control cells (n=5 independent replicates, data are mean ± SEM). (C) The ethidium transport assay described in Fig. 2B was performed with varying concentrations of ethidium bromide. The vector-subtracted median fluorescence intensities for SbMATE WT-expressing cells at the different ethidium bromide concentrations were used to calculate the transport rates, which were then normalized against the rate at 500 μM ethidium bromide set to 100 % [n=3 independent replicates, symbols are mean ± SEM, dotted line represents a hyperbolic fit, using y=P1x/(P2+x), where P1 and P2 represent the maximum observed transport rate and concentration of EtBr (x) that produces half-maximal transport rate, respectively.].

In order to further demonstrate that ethidium transport in our yeast system is SbMATE-specific, we engineered a mutation at the highly conserved aspartate residue on transmembrane helix 1, which was previously shown to be important for the protonation-deprotonation cycle crucial for substrate transport by other prokaryotic MATE family transporters. By sequence homology (Fig. 3A), the corresponding residue on SbMATE was found to be aspartate D127. We made a conservative asparagine substitution on this residue to create the mutant SbMATE D127N, and tested its ethidium transport capability. The expression levels of SbMATE D127N were found to be comparable to a specific construct of WT SbMATE protein, which was cloned with a HRV 3C protease cleavable His8 tag on the C-terminus (termed SbMATE-WT-3C-His) (Fig. 3B, 3C). The ethidium transport activity of the D127N expressing cells was found to be ~9% of those expressing SbMATE-WT-3C-His (Fig. 3D), verifying a role of D127 in organic cation transport.

**Figure 3:**
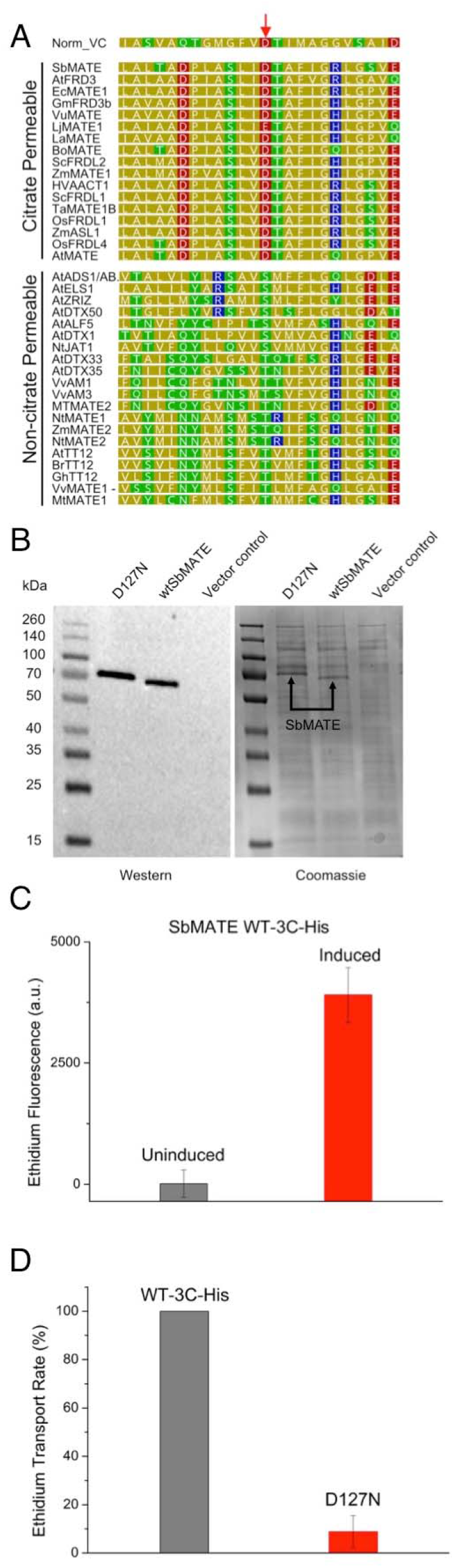
Substitution D127N impairs ethidium transport by SbMATE. (A) Sequence alignment of SbMATE with well characterized bacterial NorM from *V. Cholera* (top row) and plant MATEs, highlighting conservation of a catalytically important aspartate residue (indicated by the red arrow) across bacteria and plant MATEs associated with organic acid transport. (B) Western blot (left) and coomassie stained SDS-PAGE (right) analysis of equal amounts of *Pichia pastoris* cells expressing equivalent amounts of either SbMATE D127N, SbMATE WT, or alternatively transformed with empty vector (C) Ethidium transport was conducted as described in Fig. 2A except SbMATE WT was replaced by SbMATE WT-3C-His (see main text for details). SbMATE WT-3C-His transports ethidium efficiently (n=3 independent replicates, mean ± SEM). (D) Ethidium transport was performed as per Fig. 2A, except with induced cells transformed with SbMATE WT-3C-His, SbMATE D127N, or vector control. Transport rates were calculated as described in Fig. 2C, with the rate for SbMATE WT-3C-His set to 100 % (n=3 independent replicates, data are mean ± SEM).

These results demonstrate that SbMATE is capable of transporting both organic acid anions, like citrate, and organic cationic compounds, such as ethidium.

### Substrate transport by SbMATE can be proton and/or sodium driven

We further substantiated our results suggesting a substrate-proton antiport mechanism for SbMATE, by measuring the pH-dependence of citrate and ethidium transport. In our previous findings obtained using two-electrode voltage clamp (TEVC) to measure whole-cell currents from Xenopus oocytes, SbMATE showed greater electrogenic transport in pH 4.5 compared to pH 7.5, an effect not evident in control non-expressing cells (*8*). We proceeded to further characterize the kinetics of SbMATE transport with respect to the proton-motive force. SbMATE-mediated currents were measured at increasing extracellular H^+^ concentrations. As illustrated by the current-voltage relationships obtained at the different pH values (H^+^ concentrations ranging from 30 nM to 31 µM), SbMATE-mediated inward currents increased as the bath was acidified (Fig. 4A). The H^+^-dependent SbMATE currents at a given holding potential saturates within the experimental H^+^ concentration range, at around 30 µM H^+^ (Fig. 4B). The hyperbolic nature of these relationships suggests that only one H^+^ binds to the transporter. The Km^H^ calculated from the Michaelis-Menten relationship exhibited pronounced voltage dependence, with the apparent affinity constant exponentially increasing as the membrane potential was depolarized (Fig. 4C). These observations are consistent with the pH-dependence of the SbMATE-mediated electrogenic transport reported earlier for SbMATE expressed in X. oocytes (*8*), as well as ^14C^ citrate permeability and pH dependence reported for other plant MATEs mediating Al-exclusion in maize (*13*) and rice bean (*9*).

**Figure 4:**
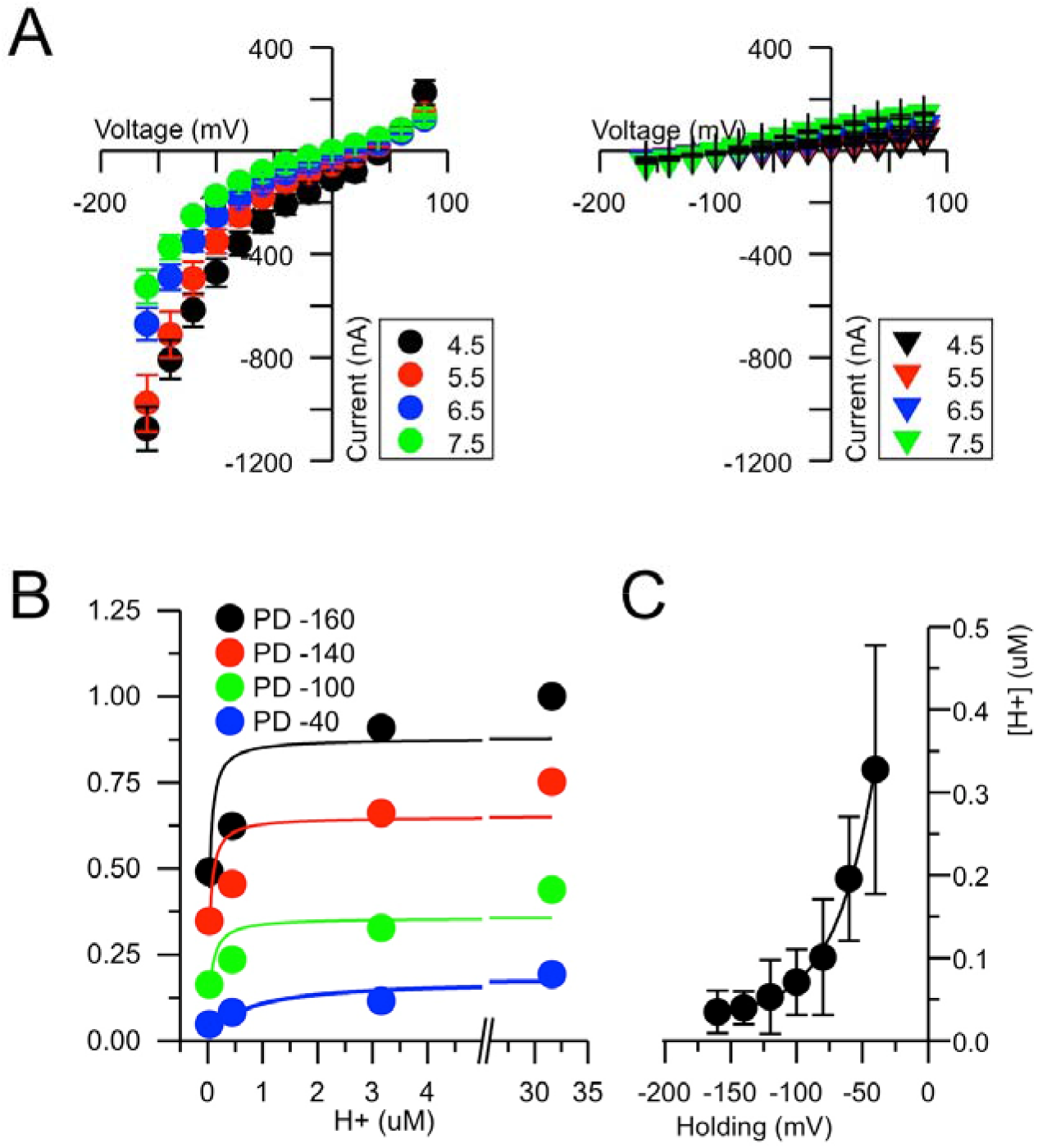
Voltage and pH dependence of SbMATE-mediated transport in X. oocytes. (A) I/V curves for currents recorded from oocytes expressing SbMATE (left panel) or control (right panel) cells in 1 mM Na^+^ bath solution at the different extracellular pHs as indicated by the different colored symbols. (B) SbMATE current magnitudes as a function of external H^+^. The K_m_^H^ values for each voltage were determined from the Michaelis-Menten function fitting of the SbMATE current magnitudes as a function of external H^+^ concentrations at a given voltage. Steady currents were normalized to the current elicited at -160 mV at pH 4.5. (C) _H_ Effect of voltage on the apparent affinity constants for protons (K_m_^H^) derived from (B). The curve shown represents a single exponential function ([H]=[H0] exp (V/τ0), with H and V being substrate and voltage, respectively, and extrapolated. Fitting parameters were H0= 1 ± 0.1 μM and τ0 = 30 ± 2 mV.

Similarly, the proton-dependence of ethidium transport into yeast cells by SbMATE was investigated by reducing the extracellular pH from 8.5 to 6.0 (i.e. thereby reducing the outwardly-directed proton gradient) during ethidium bromide incubation and subsequent washing steps. There was a notable decrease in the ethidium bromide transport rates to ~51 % (at pH 7.4) and ~33 % (at 6.0) of the rate at pH 8.5 (Fig. 5A). Since the salt concentrations were kept identical in all pH treatments, the reduction of ethidium uptake is likely a product of the decrease in the outwardly-directed proton gradient.

**Figure 5:**
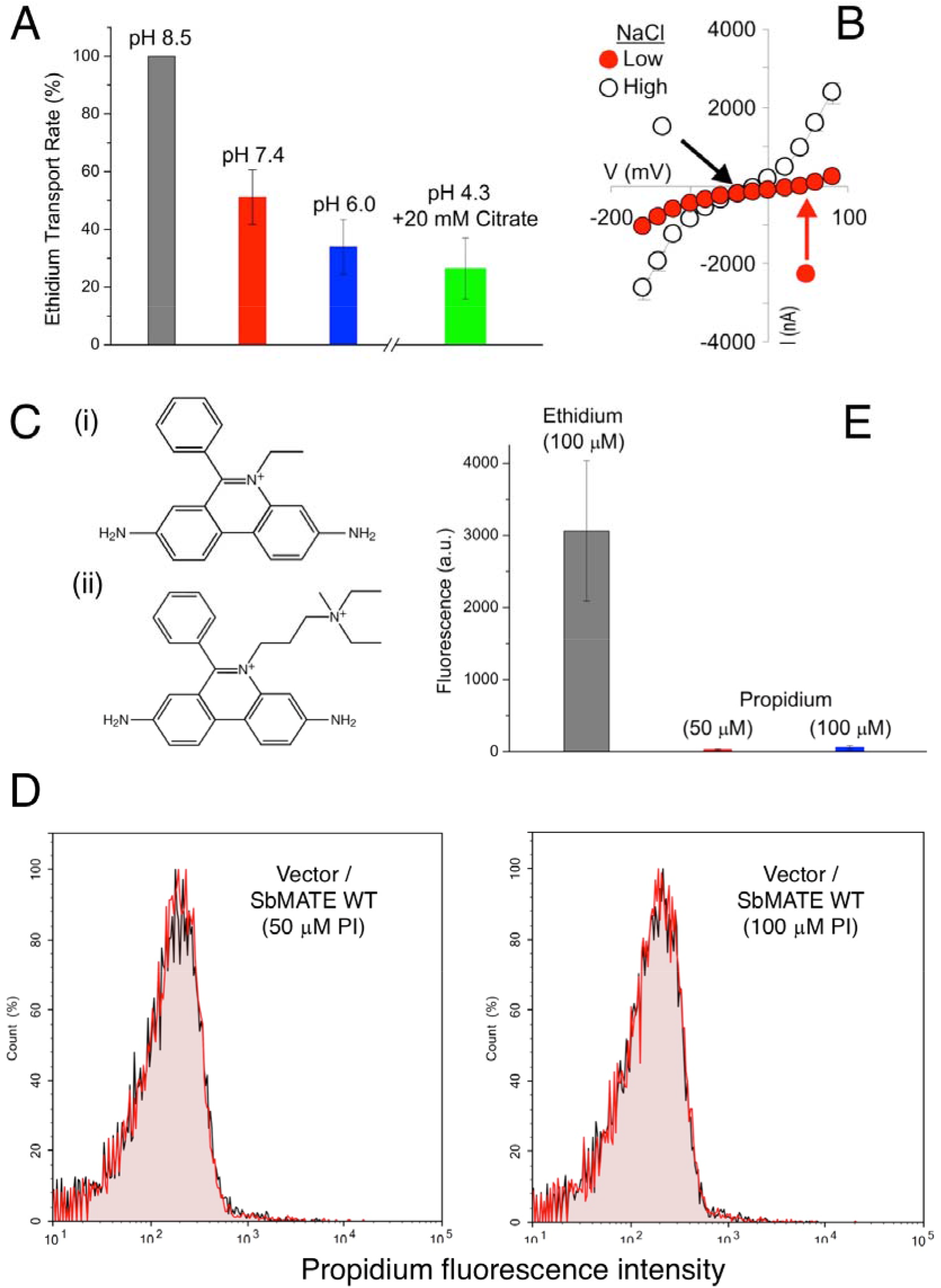
Ethidium transport by SbMATE is concentration and proton-dependent and is selective against transporting the divalent organic cation, propidium. (A) The ethidium transport assay was performed at varying extracellular pH conditions, while maintaining the same salt content as in Fig. 2B. The transport rate was calculated as described for Fig. 2C, with the rate at pH 8.5 set to 100 % (n=3 independent replicates, data are mean ± SEM). The pH 4.3 assay conditions were obtained using 20 mM citrate buffer. (B) Mean current to voltage (I/V) relationship for SbMATE-expressing oocytes recorded in different ionic (bath solution) conditions, where the reduction in NaCl in the bath media from 96 to 1 mM NaCl, pH 4.5, resulted in a decrease in the magnitude of SbMATE-mediated inward currents and a positive shift in the E_rev_ (the potential at which the net current is zero). (C) Chemical structures of (i) ethidium (+1) and (ii) propidium (+2) reveal a high degree of similarity, except for the additional charged moiety on propidium. (D) Transport assay was performed as described in Fig. 2A, except with 50 or 100 μM of propidium iodide (PI) instead of ethidium bromide. Histograms are shown for induced SbMATE WT-expressing or non-expressing control cells (n=3). (E) Graph was generated as described in Fig. 2B for induced SbMATE-expressing cells. Data from the transport assay performed with 100 μM ethidium bromide is shown for comparison. For data with 50 μM ethidium bromide, see Fig. 2C (n=3 independent replicates, data are mean ± SEM).

The cytosolic pH of yeast cells is thought to remain stable around neutrality in response to shifts in external pH between 3.0 and 7.5 (*14*). As such, we were surprised to see significant remnant transport activity by SbMATE in both Xenopus oocytes and yeast cells, even under conditions with minimal proton gradients (Figs 5A, 4A & 1B). Thus, we rationalized that SbMATE might be able to use an alternate energy source, such as the electrical membrane potential component of the proton-motive force, or another ion gradient, to power substrate transport in the absence of significant proton gradients. Consistent with this hypothesis, in Xenopus oocyte whole-cell recordings, reducing [Na^+^] in the bath media by lowering the NaCl concentration from 96 to 1 mM NaCl, thereby reducing the inwardly directed Na^+^ gradient, resulted in a reduction of the SbMATE-mediated inward and outward currents and a shift in E_rev_ to positive potentials (Fig. 5B).

Overall, the above results indicate that SbMATE mediates citrate and ethidium transport coupled to high-affinity H^+^ binding and/or potentially other cations, such as, Na^+^, validating a proton (H^+^)/sodium (Na^+^)-substrate antiport mechanism.

### Substrate and charge selectivity of SbMATE

Citrate is a physiological transport substrate of SbMATE. Since citrate buffers are used to maintain acidic pH ranges, we challenged SbMATE-expressing yeast cells with an external pH of 4.3 and measured the uptake of 50 µM ethidium bromide, in the presence of 20 mM citrate and the same salt concentrations used in all of the above-described yeast experiments (i.e. 137 mM NaCl, 2.7 mM KCl). Amidst an almost complete lack of an outwardly-directed proton gradient, and despite an inwardly-directed sodium gradient, SbMATE still showed an appreciable degree of ethidium uptake (Fig. 5A). The transport rate was reduced to 25% of that observed at pH 8.5 (Fig. 5A), which is slightly lower than the 33% transport rate observed at pH 6.0 (Fig. 5A), and still above the background of nonexpressing cells. These results suggest that the transport of organic cations by SbMATE can occur even in the background of highly competing levels of citrate, or other organic acid anions.

Similar to ethidium, its structural and functional analog, propidium, also binds to nucleic acids and exhibits an increase in fluorescence. However, in contrast to monovalent ethidium, propidium is a divalent organic cationic dye (Fig. 5C), which serves as a test compound to study the substrate charge preferences of SbMATE. Conducting the yeast transport assay with 50 μM and 100 μM propidium iodide, SbMATE-mediated transport was found to be negligibly low, compared to the transport of ethidium at the same concentrations (Fig. 5D, E). This observation suggests that SbMATE can distinguish between monovalent vs divalent organic cationic compounds, preferentially transporting the former. Propidium iodide is also routinely used to assess the viability of cells in flow cytometry experiments. Thus, the low fluorescence intensity of SbMATE-expressing cells with propidium (Fig. 5D, E) also demonstrates that our cell sample preparations were intact and healthy. As such, the increase in ethidium fluorescence in our experiments (Figs. 2A, 3C) was most likely due to transport, as opposed to alterations in overall membrane permeability.

### Purification of functionally-folded SbMATE

Following the confirmation that SbMATE over-expressed in *Pichia pastoris* was transport-competent (Figs. 2, 3 and 5), we proceeded with purifying the protein in detergent solution from 8 L fermenter cultures. Briefly, bDDM detergent-solubilised SbMATE was purified by affinity chromatography using the His_10_ tag and size-exclusion chromatography. Each 8 L fermenter run produced an average yield of ~5 mg of purified and monodisperse SbMATE protein (Fig. 6A).

**Figure 6:**
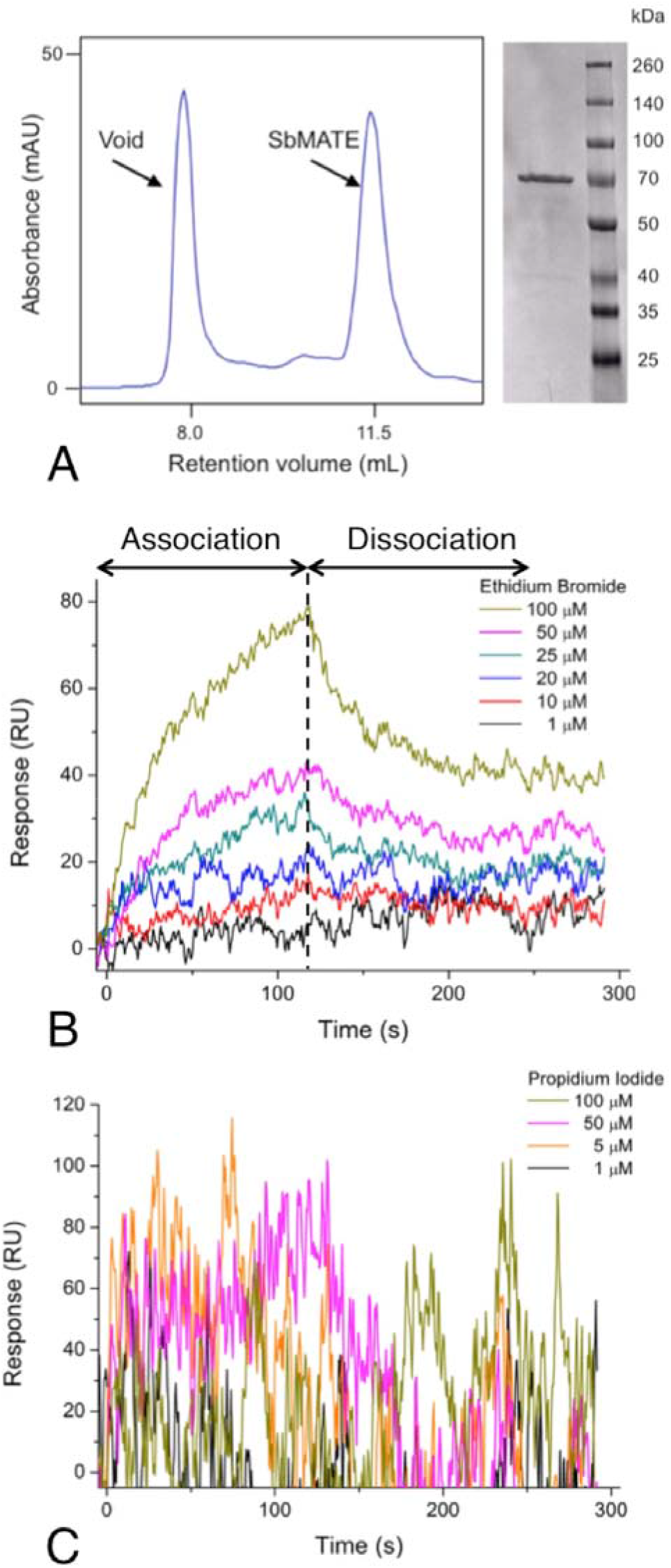
Purified SbMATE binds ethidium bromide, but not propidium iodide. (A) Detergent purified SbMATE elutes from size exclusion chromatography as a symmetrical monodispersed peak (FPLC gel filtration trace on left) that is deemed to be ~95% homogeneous (coomassie stained SDS-PAGE of pooled FPLC peak fractions shown on right). (B) Purified SbMATE was covalently immobilized on a CMD surface plasmon resonance (SPR) sensorchip. 100 μL of varying concentrations of ethidium bromide were injected, with each concentration having an association and dissociation time of 120 s. The chip was regenerated between each injection using a short 10 μL injection of 50 mM glycine-HCl, pH 2.1. Traces on the sensorgram represent double-referenced data, obtained by subtracting response signals from a blank surface and buffer injections (n=5 replicates performed using independently purified batches of protein, traces represent data from a single experiment). (C) SPR was conducted as described in Fig. 6B, except with varying concentrations of propidium iodide instead of ethidium bromide (n=3 independent replicates, sensorgrams are from a single experiment).

The functional integrity of detergent-purified SbMATE protein was tested using two different approaches. First, we conducted an *in vitro* binding assay using label-free, surface plasmon resonance (SPR) to measure the binding kinetics of SbMATE to the transport substrate, ethidium. Purified SbMATE was observed to bind ethidium in a saturable manner (Fig. 6B), with a dissociation constant (*K*D) of 47.1 ± 15.6 μM (mean ± SEM, n=5) (Fig. 6B). This binding affinity is relatively high compared to the known SbMATE substrate, citrate, which has been tested at higher μM-mM concentrations. Consistent with our transport assays (Fig. 5D, E), we also found that the use of propidium iodide, at the same concentrations as ethidium in this assay, did not result in discernible, concentration-dependent binding traces (Fig. 6C).

Functional validation of detergent-purified SbMATE was further performed by reconstituting the protein into an artificial membrane environment and evaluating its electrogenic activity (i.e. the capacity to transport ions), using electrophysiological measurements. Incorporation of SbMATE resulted in single channel-type electrogenic activity, seen as discrete changes in current amplitude (with the unitary channel spending about half of the total recording time in a conducting open state of about 1 pA) as the transport protein transitioned between open and closed conformational states (Fig. 7). This type of single channel activity, is consistent with that recently reported for other plant MATE transporters overexpressed in *Arabidopsis* (*15*).

**Figure 7:**
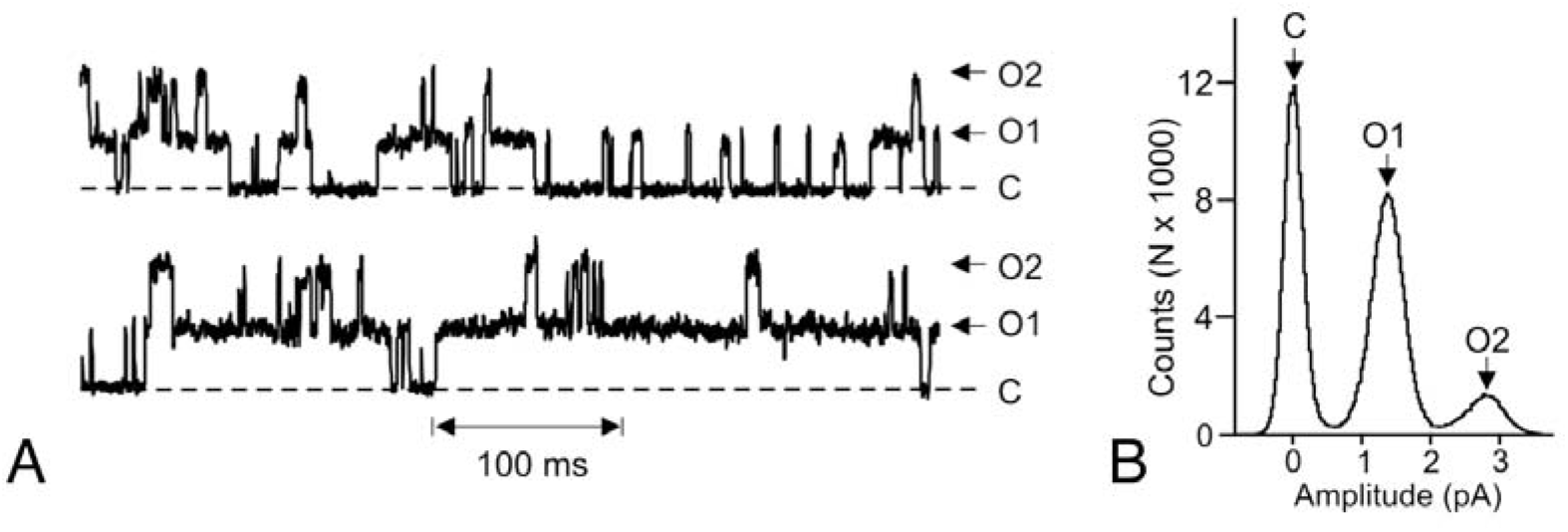
Functional reconstitution of detergent purified SbMATE. Representative recordings for the purified SbMATE recombinant protein expressed in *P. pastoris* reconstituted into liposomes and incorporated into lipid bilayers in an asymmetric ionic *cis:trans* gradient (200: 20 mM NaCl, pH 7.5). (A) Representative current recording following addition of SbMATE containing proteolipomes to the *cis* side of the artificial lipid bilayer. O and arrows indicate open states. C and dashed lines indicate the zero current closed state. Time scale is denoted on the bottom margin. The zero current level (closed state) in all cases is indicated by the horizontal dotted lines. (B) All point histograms illustrating the open and close state distributions for the full length recordings illustrated in (A). The zero current level (closed state) and various open states are above each peak

### High-affinity nanobodies against purified SbMATE

The availability of large amounts of functional purified protein is valuable for the discovery and screening of potential binders, such as antibodies, which can be useful as research reagent tools. Integral membrane proteins, however, are typically considered ‘hard targets’ for discovering antibodies *via* the traditional animal immunization approach. Additionally, plant-derived proteins, in general, are known to have substantially lower immunogenicity compared to other microbial/mammalian antigens that are used to immunize animals. To surpass these hurdles, we applied an *in vitro* approach (*16*) to enable the rapid discovery of single-domain antibodies, also called nanobodies (Nbs), against membrane protein targets.

Our Nb discovery process resulted in a panel of six Nbs (Table 1) that were purified from *E. coli* and tested for binding to SbMATE. The Nb panel was diverse in binding affinities, ranging from low-affinity (μM) (Fig. 8A-D) to moderately high-affinity (nM) (Fig. 8E, F), as characterized by SPR. The higher affinity binders, Nbs 86 and 94, were further characterized to have dissociation constants, *K*_D_ of 173.57 ± 45.53 nM and 99.53 ± 19.32 nM, respectively (mean ± SEM, n=3) (Fig. 8E, F). These represent novel, first-generation Nbs obtained from a *naïve* camelid-derived Nb library, without conducting deliberate affinity maturation.

**Figure 8:**
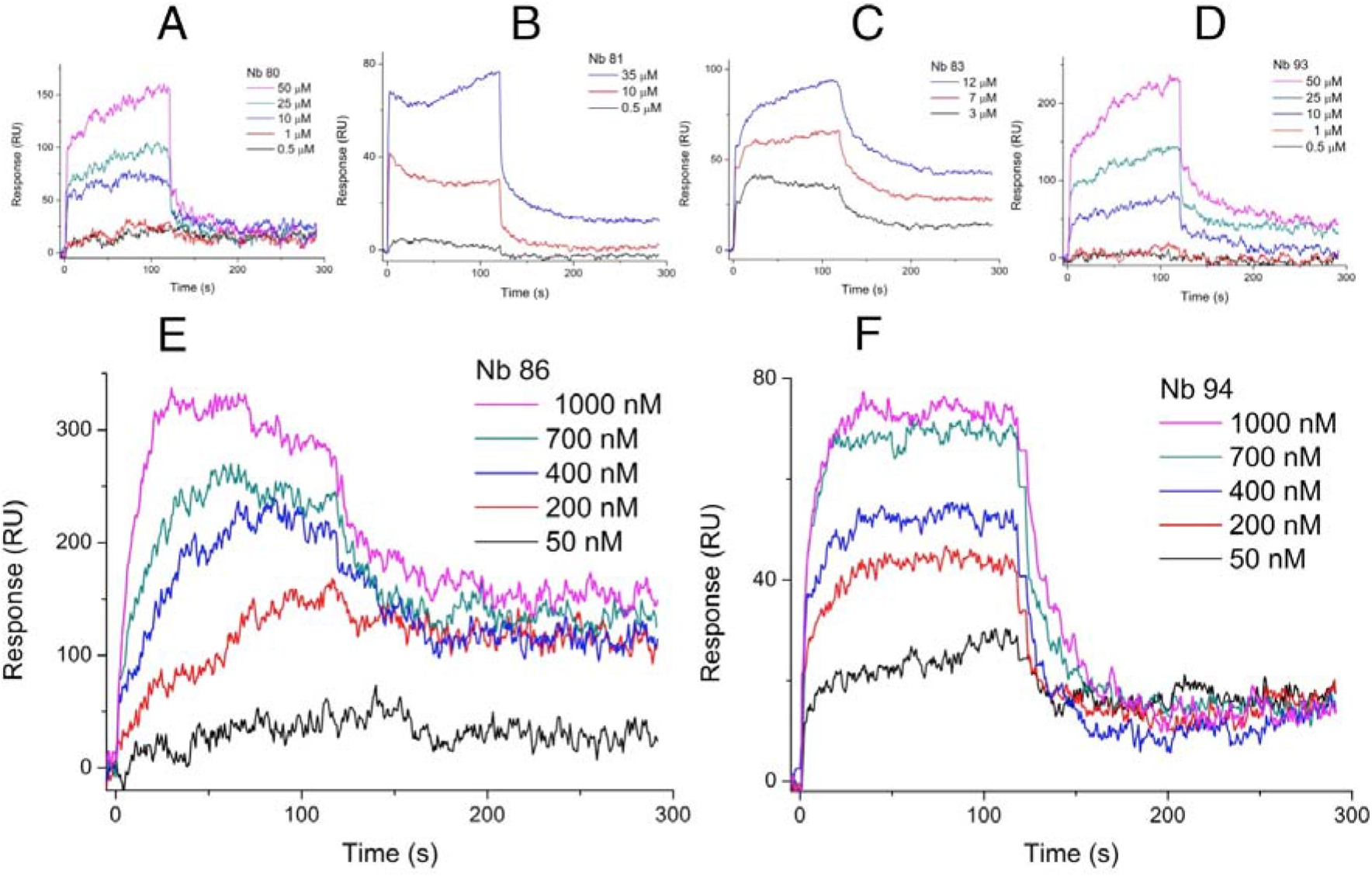
*In vitro* discovered nanobodies bind to purified SbMATE. SPR was performed and the data were double-reference subtracted as described in Fig. 6B, except purified nanobodies (Nbs) were injected at various concentrations, instead of ethidium. The calculated dissociation constants (*K*D) for the lower-affinity Nbs were (A) Nb 80 – 12.2 μM, (B) Nb 81 – 23.9 μM, (C) Nb 83 – 3.3 μM, and (D), Nb 93 – 26.1 μM. (E) Nb 86 and (F) Nb 94, bound with higher affinity, with *K*D’s in the nM range. To get more accurate *K*D values, a BSA-immobilized surface was used as the reference channel, instead of a blank surface. Calculated *K*D’s; Nb 86 – 173.57 ± 45.53 nM; Nb 94 – 99.53 ± 19.32 nM (n=3 independent replicates, sensorgrams are from a single experiment).

**Table 1:**
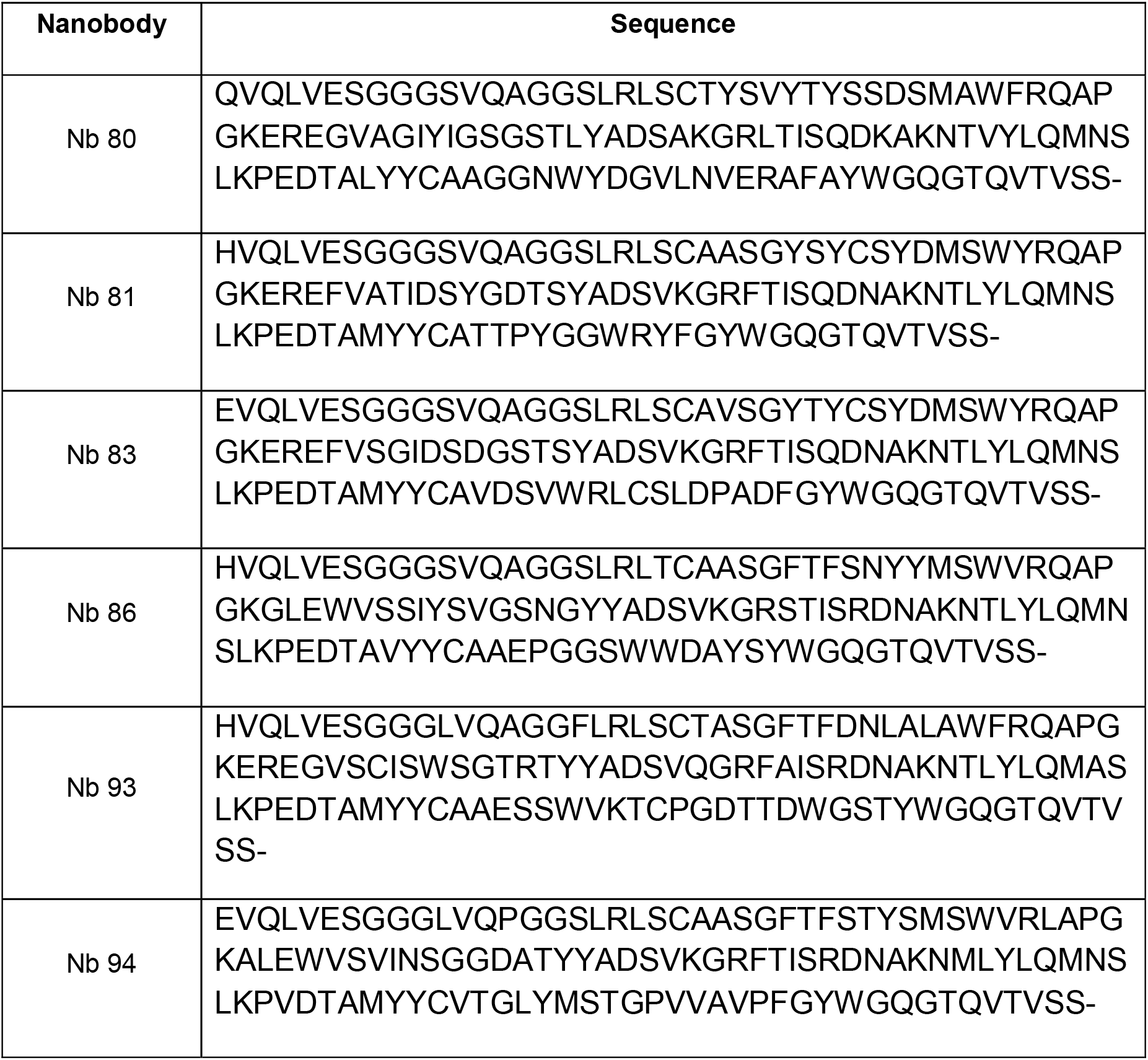
Nb protein sequences. All Nbs have the expression vector-added leader sequence, MRGSHHHHHHGMASMTGGQQMGRDLYDDDDKDHPFT

### Nanobodies bind to the folded conformation of SbMATE

One advantage of *in vitro* Nb technologies is that discovery of Nbs occurs against the full-length, natively folded target proteins. Our purified SbMATE protein was functionally folded, as was demonstrated by its ability to bind the transport substrate ethidium, to distinguish the non-transported analog, propidium, and also its ability to mediate ion transport when reconstituted into an artificial lipid bilayer (Fig. 6B, C and Fig. 7). In order to investigate whether some of our Nbs bind to a native fold of the antigen, we tested the effect that Nb binding might have on ethidium binding to SbMATE and on SbMATE-mediated electrogenic transport.

When we performed SPR using Nbs and ethidium at a molar ratio 1:10 Nb:ethidium, our two high-affinity Nbs, 86 and 94, reduced ethidium binding on SbMATE to 35.04 ± 13.39 % and 71.40 ± 13.91 %, respectively (mean ± SEM, n=3) (Fig. 9A, B). In contrast, these Nbs did not yield a reduction of the non-specific ethidium signals obtained for the hydrophobic control protein, BSA (Fig. 9C). Furthermore, upon extracellular exposure of SbMATE-expressing Xenopus oocytes to a mixture of Nbs (Nbs 83, 86, 89, 94 and 124), a ~2-fold increase in the SbMATE mediated inward currents were observed. This agonistic effect of the Nbs mix was reversible upon re-establishing the control bath conditions (Fig 10A). Likewise, intracellular Nb-SbMATE interactions were studied in SbMATE expressing cells preloaded with the Nb mix 2.5 hours prior to the electrophysiological recordings. This resulted in larger currents than control SbMATE-expressing cells preloaded with water (Fig 10B). Importantly, neither extracellular or intracellular Nbs had any effect of the current magnitudes recorded in non-expressing cells. These findings are interesting as they indicate an agonistic effect for our Nbs on SbMATE’s transport functions.

These findings suggest that our *in vitro-*discovered Nbs bind to the transport-competent, folded form of SbMATE, thereby highlighting the suitability and advantages of such *in vitro* Nb discovery technologies when studying integral membrane proteins, including those from plants.

**Figure 9:**
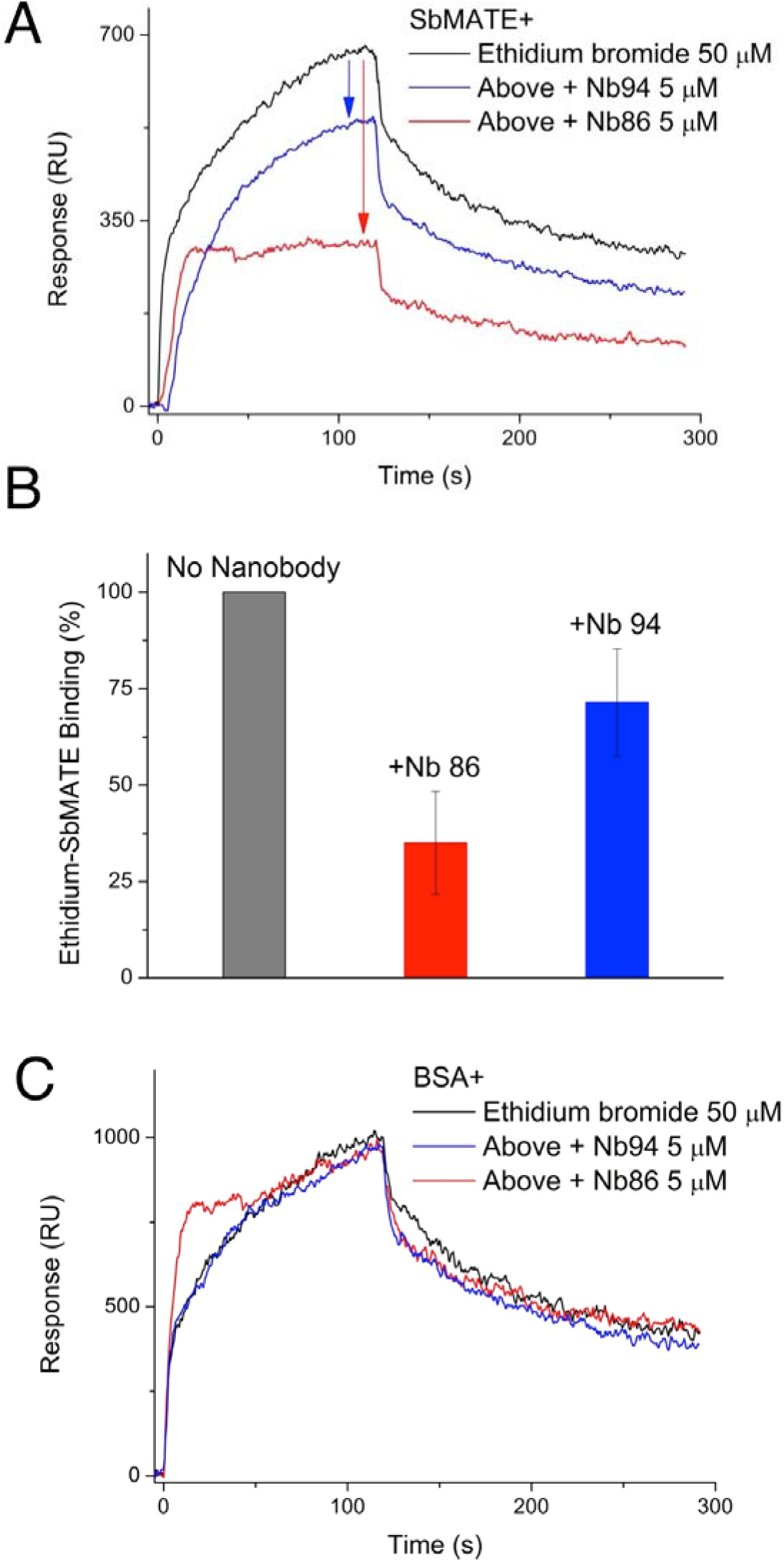
High-affinity Nbs inhibit ethidium binding to SbMATE. (A) SPR was performed as described in Fig. 6B, with the following modifications. 50 μM ethidium bromide was injected either directly on SbMATE alone, or following injections of 5 μM Nb 86 or Nb 94 without a subsequent regeneration step. The ethidium trace (top/black) represents double-referenced data as described in Fig. 6B, while the ethidium + Nb traces (Nb 86 – bottom/red; Nb 94 – middle/blue) were generated using a similar double-referencing method, except the respective Nb injections alone were used instead of a buffer injection for the blank subtraction (n=3 independent replicates, sensorgram is from a single experiment). (B) The response value (in RUs) at the end of the association phase (i.e. at ~118 s) of the injections described in Fig. 9A were normalized, with the RU value for the ethidium-SbMATE trace set to 100%. Response RUs were reduced to 35.04 ± 13.39 % for ethidium-Nb86-SbMATE and to 71.40 ± 13.91 % for ethidium-Nb94-SbMATE (n=3 independent replicates, mean ± SEM). (C) SPR was performed as described in Fig. 9A, except the non-specific hydrophobic control protein, BSA was immobilized on the sensor chip, instead of SbMATE. To obtain comparable ethidium signals, 2x higher levels of BSA than SbMATE were immobilized.

## DISCUSSION

Worldwide food demand is projected to experience a 40% increase by 2030 (*17*). Studies on crop yields predict that this is unlikely to be met by applying traditional agricultural methods based on conventional plant breeding and additional use of fertilizer and chemicals. Novel technologies will instead need to focus directly on improving various agronomic properties of crops. For example, a molecular breeding approach based on the use of markers linked to *SbMATE* was used to improve sorghum Al tolerance and yields on acid soils (*18*) and biotechnological approaches were applied to improve Al-tolerance in barley, via transformation with the *SbMATE* gene (*19*). A detailed understanding of substrate specificities and transport mechanisms of agronomically important transporters, such as SbMATE, will provide new avenues for engineering more effective (Al-tolerant) SbMATE-type transporter proteins and aid the success of these approaches.

To date, the characterization of the subgroup of plant MATE transporters associated with Al resistance and Fe transport/homeostasis has been solely performed in relation to their organic anion transport activity (specifically citrate) which underlies their major *in planta* roles. MATE family proteins however, were initially identified as proteins that could transport a wide range of organic cationic substrates. In fact, in plants, a separate clade of plant MATEs have been associated with the transport of a broad variety of secondary metabolites (Fig. 1A; also see (*2*)). Our findings, indicating that SbMATE can also transport organic cationic compounds, such as ethidium, brings these clades closer together under the larger umbrella of MATE family transporters. It is interesting, however, that propidium, which is a bivalent structural and functional analog of ethidium, was not transported as efficiently (Fig. 5). Although propidium’s mass is slightly larger than that of ethidium, substrates of MATE family transporters generally have a wide range of molecular weights (*20*). It is thus likely that the lack of propidium transport by SbMATE suggests an inherent limitation on the number of positive charges that can be accommodated within SbMATE’s substrate-binding site(s), and is not simply an effect of differential mass given the variable size of MATE-family substrates.

Our mechanistic studies on SbMATE strongly suggest that its transport cycle is driven by proton and/or sodium gradients, as was recently shown for the bacterial MATE transporter, NorM (*21*). The ethidium transport activity of SbMATE in yeast was greatly reduced (to 33%) but not abolished, when the extracellular pH was lowered from pH 8.5 to 6.0 (Fig. 5A). The residual transport activity at pH 6.0 indicates a markedly low p*K*a for protonation sites, such as D127, on SbMATE compared to other known MATE transporters. This is not surprising, given that SbMATE is activated in sorghum crops growing in acidic soils at pH <5.0.

Perhaps most interesting is our finding that ethidium uptake continues to occur even in the presence of high levels of the physiological substrate, citrate, and amidst an almost nonexistent outwardly-directed proton gradient (pH_out_ of 4.5), and substantial inwardly-directed sodium gradient at 157 mM Na^+^_out_. First, it is unclear as to what exactly drives this residual ethidium transport under these conditions, and bears some similarity to the transporter-mediated facilitated downhill fluxes observed in the absence of proton-motive forces for secondary transporters of the Major Facilitator Superfamily (MFS) (*22*). Secondly, owing to dedicated yeast carboxylate transporters, citrate is generally bioavailable to yeasts such as *P. pastoris* in culture medium and should be available for transport via the membrane-embedded SbMATE protein (*23–25*). Thus, our findings implicate a strong physiological relevance for organic cationic substrates of SbMATE that might compete with citrate in acidic soils.

The SbMATE residue D127 is highly conserved as a negatively charged residue throughout bacteria and all plant MATE transporters associated with organic acid transport, while being highly conserved as a neutral (serine or threonine) residue in plant MATE transporters associated with the transport of other plant metabolites (Fig. 3A). In the bacterial MATE transporters, the corresponding carboxylate residue is D32 in NorM from *Vibrio parahaemolyticus*, D36 in NorM from *Vibrio cholerae*, D41 in NorM from *Neisseria gonorrheae*, D41 in the PfMATE transporter from *Pyrococcus furiosus*, and D40 in DinF-BH from *Bacillus halodurans*. These residues have been shown to be crucial for transport activity, and are likely involved in proton/sodium and/or ion-substrate competition. Consistent with these findings, our protonation-mimetic mutation, D127N, severely impaired SbMATE’s ability to transport ethidium in yeast (Fig. 3). Thus, the eukaryotic SbMATE transporter appears to show remarkable structure-function homology with bacterial MATE transporters, even with a considerably extended intracellular N-terminal region that is absent in the bacterial MATE transporters studied to date.

*In vitro*, functional biochemistry and structural biology of plant-derived integral membrane proteins have been hampered due to the lack of reliable, large-scale recombinant protein production systems. Our *P. pastoris* yeast system is capable of yielding milligram levels of purified SbMATE protein, which was found to be functional in terms of binding to the transport substrate, ethidium, but not to propidium (Fig. 6). The channel-like current activity observed in our electrophysiological experiments extends recent reports of other active transporters exhibiting similar currents (*15, 26*). These protein preparations can be used to screen additional transport substrates, and perhaps even transport activators that could improve Al-tolerance phenotypes.

Our panel of SbMATE-binding Nbs (Fig. 8), including some that compete with ethidium, display agonistic effects on the SbMATE-mediated electrogenic transport in oocytes (Fig. 10), when administered extra-or intracellularly. These results suggest that one or more Nb-binding epitope(s) on SbMATE are accessible from either side of the membrane, such as the substrate-binding sites in the central cavity of the transporter. As such, the agonism observed could possibly be a result of Nb-binding induced proton/sodium transport, similar to substrate-induced proton transport seen in NorM, PmpM, DinF, and PfMATE (*21, 27-29*). Such panels of Nbs are useful, not only as tools to study protein trafficking under various environmental conditions, but to also potentially alter transporter efficiencies, and therefore, their functional roles *in planta.*

**Figure 10:**
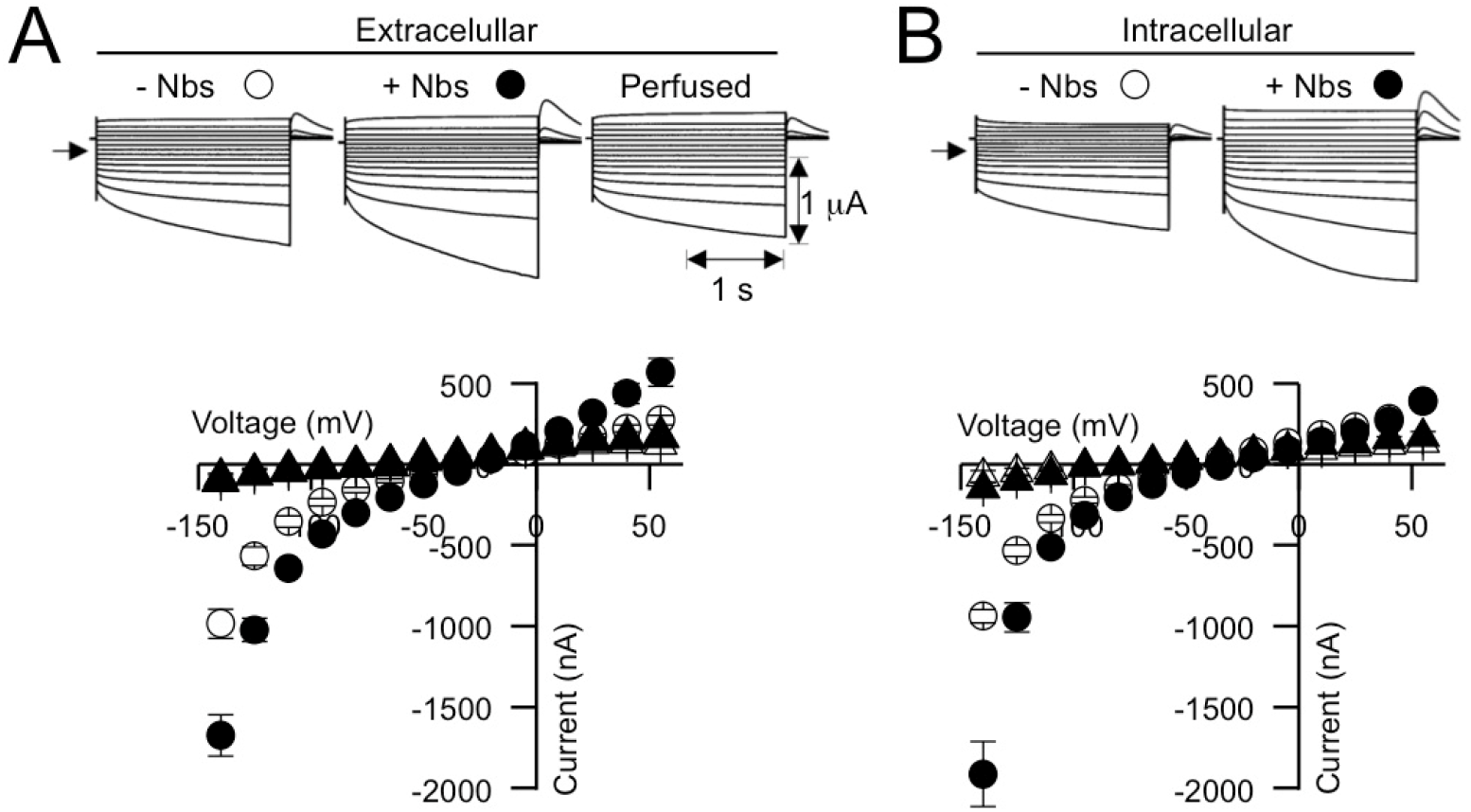
High-affinity Nbs modulate SbMATE-mediated transport activity in X.oocytes. Representative currents recorded under voltage clamp and corresponding mean current to voltage (I/V) relationships from oocytes expressing SbMATE. The symbols for each I/V curve correspond to the symbols depicted at the top of each set of traces above the I/V. Currents were recorded in control cells (not expressing SbMATE) under identical sets of conditions (traces not shown) and are depicted by the white (-Nbs) and black (+Nbs) circles. (A) Currents were recorded in a bath solution consisting of a Ringer solution (in mM: 96 NaCl, 1.8 CaCl_2_, 1 KCL) (pH 7.5) prior to (-Nbs) and following perfusion with the Ringer solution containing a mixture of Nbs (+Nbs). The mixture of Nbs consisted of Nb*X* (final concentration in μM), *x* being 83 (14), 86 (3), 89 (1.4), 94 (0.7) and 124 (0.9). The third set of traces labeled “perfused” illustrates representative currents recorded following removal of the Nbs mixture by perfusing for 3 min with the control (-Nbs) solution. The holding potential in each was set to 0 mV and voltage pulses were stepped between -55 and -140 mV (in 15 mV increments). The arrow on the left of the set of traces represents the zero current level. Current and time scales are given at the bottom right of the set of traces. (B) Currents were recorded in a bath solution consisting of the same Ringer solution described in A. Recordings were in SbMATE-expressing oocytes that were preloaded with a Nb mixture or water at 2.5 h prior to the recording. Preloading was achieved by injecting each cell with 48 nL of water or a 100 fold Nbs mix used in A.

## CONCLUSION

In this study, we have used a combination of Xenopus oocytes and a yeast-based heterologous recombinant expression system to functionally characterize the transporter protein, SbMATE from *Sorghum bicolor*, which is responsible for imparting Al-tolerance in sorghum. In doing so, we have extended the substrate recognition profile of SbMATE, demonstrating a H^+^/Na^+^ coupling transport system with the ability to not only transport citrate, but also organic monovalent cations, such as ethidium. We have also shown that yeast-expressed SbMATE can be purified in large quantities, and that the purified protein is functionally-folded and amenable as an antigen for *in vitro* antibody discovery. The overall Nb discovery methodology is also applicable to other plant-derived membrane proteins, allowing for the rapid development of reagent tools to probe protein localization and function.

## METHODS AND MATERIALS

### Expression and Functional characterization of SbMATE in *Xenopus* oocytes

cRNAs were prepared using the mMessage mMachine T7 *in vitro* transcription kit (Ambion, http://www.ambion.com/) with 1 μg of *ScaI*-linearized T7TS plasmid template containing the SbMATE coding region, flanked by the 3′- and 5′-untranslated regions of a Xenopus β-globin gene. *Xenopus laevis* oocyte harvesting, microinjection, and whole-cell recordings under two electrode voltage clamp methods were performed as described previously (*30*). Briefly, V–VI stage oocytes were injected with 48 nL of water containing 20 ng of cRNA encoding SbMATE. ^14C^Citrate efflux control and SbMATE-expressing oocytes (2 days after cRNA injection) were injected with 23 nL of 460 μM ^14C^citrate (1 nCi/oocyte). The cells (6 cells per replicate; n=4) were allowed to recover for 3 min in ice-cold Ringer solution (pH 4.5) and then transferred into 1.5 mL Ringer solution (pH 4.5) at room temperature. At the indicated time points, 0.75 mL of the bathing solution was sampled and replaced with fresh buffer. At the end of each experiment the cells were disrupted in scintillation fluid. Radioactivity from sampled efflux buffer at the various time points and remaining radioactivity in the oocytes was counted using full-spectrum DPM counting in a Beckman Coulter LS6500 Liquid Scintillation counter. For two-electrode voltage clamp, whole-cell currents were recorded 2–3 days following the microinjections using an Geneclamp 500B amplifier/PClamp 10 data acquisition system (Molecular Devices, Axon instruments). Recordings were performed under voltage clamp in various bath solutions containing 1 mM KCl and 1.8 mM CaCl_2_ in the background. Na^+^ was supplemented as NaCl (up to 96 mM), and the pH of the various solutions was adjusted to between 4.5 and 7.5 (as indicated in the figure legends). The osmolarity or all solutions was adjusted to 220 mosmol kg^-1^ using D-sorbitol. The current-voltage (I/V) relationships were constructed by measuring the current amplitude at the end of the test pulses. All recordings were performed in four biological replicates (i.e. two frog donors). Error bar represent SEM and are not shown when smaller than the symbol.

### Expression and Functional Characterization of SbMATE in *Pichia pastoris*

The *Sorghum bicolor* MATE gene (*SbMATE*, GenBank EF611342) was cloned into a pPICZ-C vector that was modified to incorporate N- and C-terminal FLAG and His_10_ affinity tags, respectively, or alternatively cloned with a HRV 3C protease cleavable His_8_ tag on the C-terminus (termed SbMATE-WT-3C-His). The resultant construct was linearized using *PmeI* and transformed by electroporation into KM71H cells. Transformants were plated onto YPDS agar plates (containing an increasing amount of Zeocin antibiotic; 200-1000 ug mL^-1^) and colonies screened for SbMATE expression via small scale expression and western blot analysis. High expressing strains were selected for fermentation in 8 L cultures using a Bioflow 415 bioreactor. SbMATE expression was induced by slow methanol addition (3.6 ml per hour per liter of culture volume) for 16-18 hours. The SbMATE D127N mutant was generated using the Infusion technique (Takara Bio) and expressed as above.

*P. pastoris* cells (expressing SbMATE, mutants, or the non-expressing vector control) were washed with standard phosphate buffered saline (PBS), pH 7.4, and diluted to an OD_600_ of 15 per 1 mL of experimental sample. The assay buffer for experiments conducted at pH 8.5 was 25 mM Tris-Cl, pH 8.5, 137 mM NaCl, and 2.7 mM KCl. Buffers for the different pH conditions were: pH 7.4 - PBS; pH 6.0 - 50 mM MES; pH 6.0, 137 mM NaCl, 2.7 mM KCl; pH 4.5 - 20 mM sodium citrate; pH 4.3, 137 mM NaCl, 2.7 mM KCl. Drugs, ethidium bromide or propidium iodide were incubated with cell suspensions for 20 min at RT, followed by washes in respective ice-cold buffer(s) and measurements by flow cytometry (Novocyte, Acea Bioscience). Approximately 12,000 events were collected per sample and the appropriately gated data were displayed qualitatively as histograms, and quantified through median fluorescent intensities. Additional experimental details are available in the figure legends.

### Purification of SbMATE from *Pichia pastoris*

Cells were lysed at 40 Kpsi by a single pass through a constant cell disrupter. Cellular debris was removed by centrifugation (12,500 x *g*, 20 minutes, 277K) and crude membranes prepared by centrifugation at 38,000 x g for 2-3 hours at 277K. Expressed SbMATE was bDDM detergent-solubilised from membranes and purified by affinity chromatography using the histidine tag and subsequent size-exclusion chromatography.

### Surface Plasmon Resonance Binding Studies

SPR measurements were performed on the BiOptix 404Pi instrument using CMD sensorchips. Detergent-purified SbMATE was covalently immobilized using EDC-NHS chemistry to 5, 000 resonance units (RUs) per sensor channel in a buffer containing 10 mM MES pH 6.0 and 0.05% bDDM. The running buffer was switched to 50 mM HEPES, pH 8.0, 150 mM NaCl, and 0.03% bDDM for the binding experiments with ethidium bromide, propidium iodide, or Nbs. The drug solutions were prepared and Nbs were desalted into the running buffer to minimize buffer-related effects on the SPR traces. Dissociations were performed using short injection pulses of 50 mM glycine, pH 2.1. All data were analyzed using the Scrubber (BioLogic) software, and subjected to the software-associated double-subtraction processing, curve-fitting, and fit statistics. Additional experimental details are available in the figure legends.

### *In vitro* Nanobody Generation

A detailed protocol, including reagents and oligo designs for conducting *in vitro* discovery of single-domain Nbs, is available in (*16*). Briefly, a *naïve* camelid single-domain library was purchased from Creative Biolabs and used for Nb discovery by applying an mRNA/cDNA display methodology developed in our laboratory (*16*). PCR on the phagemid was used to amplify the Nb library and add the necessary 5’ sequences for *in vitro* transcription, translation, and Nb detection. The PCR product was *in vitro* transcribed to mRNA, and ligated to a DNA linker modified with a puromycin group on the 3’ end, using a DNA splint-assisted ligation reaction. This was followed by *in vitro* translation using the PURExpress kit (New England Biolabs) and mRNA-Nb fusion formation was carried out by adding 2.5 μL of 1 M MgCl_2_ and 12.5 μL of 2 M KCl to the 25 μL reaction followed by overnight incubation at --20°C. Fusion molecules were purified using the polyA sequence on the puromycin linker and first strand cDNA synthesis was conducted using Superscript IV (ThermoFisher). mRNA/cDNA-Nb fusions were applied to negative HIS-tagged antigens to remove non-specific binders. The unbound pool of fusions was added to purified HIS-tagged SbMATE immobilized on Ni-NTA agarose magnetic beads. Both negative and positive selection incubations were for 20 min at RT on a tube rotator. The positive selection beads were washed 10 times in 1 mL. Finally, the Nb fusions were eluted from the beads and the sequence recovered using PCR. For the selection step, the buffers used were as follows: Ni-NTA Antigen-Binding buffer (20 mM Tris-HCl, pH 8.0, 300 mM NaCl, 0.05% bDDM); Nb-Ag Interaction buffer (20 mM Tris-HCl, pH 8.0, 150 mM NaCl, 0.05% bDDM, 1 ug/mL tRNA, 1 ug/mL BSA); Wash buffer (Interaction buffer with 20 mM imidazole), Elution buffer (20 mM Tris-HCl, pH 8.0, 0.05% bDDM, 300 mM imidazole). After 3 rounds of panning, the PCR product was bulk-cloned into TOP 10 cells (Invitrogen) and 96 individual clones were sequenced. Enriched sequences were cloned into pET100-D-TOPO (ThermoFisher) for expression and purification. The reader is referred to (*16*) for a detailed protocol of this process.

### Expression and Purification of Nanobodies

Expression and purification of nanobody binders was performed as previously described (*16*) with minor modifications. Briefly, pET100-D-TOPO expression vectors harboring the nanobody sequences were transformed into either BL21 (DE3) Star (Nbs 83, 94) or Shuffle T7 cells (Nbs 86, 89, 124). Large-scale growths were performed in either Terrific or Luria Broth in 1 L shaker flasks that were grown at 37ºC to ~0.7 OD_600nm_. At this time, the growth temperature was reduced to 30ºC and the cultures induced with 1 mM IPTG for 3-4 hours. The cells were then harvested (12,500 x g, 4ºC, 20 minutes) and resuspended in lysis buffer (20 mM Tris-HCl pH 8.0, 100 mM NaCl and 15% glycerol) before disruption with a constant cell disruptor (Constant Systems) at 15 Kpsi. Cell debris was removed by centrifugation (38,000 x g, 4ºC, 30 minutes) and the supernatant was collected and incubated with nickel affinity resin (Ni-NTA Agarose, Qiagen) at 4ºC for ~1 hour after the addition of 20 mM imidazole to the supernatant. The resin was then loaded into a gravity column and washed with Buffer A (20 mM Tris pH 8.0, 400 mM NaCl, 0.05% Tween-20, 20 mM Imidazole), followed by separate washes with Buffer A containing 40 mM imidazole, 1 M NaCl and 60 mM imidazole. Finally, bound protein was eluted from the resin with Buffer A containing 300 mM imidazole and concentrated to ~500 uL before a final polishing step using a Superdex 200 10/300 size exclusion column.

### Reconstitution of SbMATE transport activity in artificial lipid bilayers

The purified SbMATE protein was reconstituted into phosphatidyl choline (PC: Avanti Polar Lipids) liposomes. PC was dried under nitrogen gas to generate a thin film prior to hydration. Hydration of the dry lipid film was accomplished by adding PBS with gently agitation at root temperature, reaching a final lipid concentration of 33 mg/ml. The multilamellar vesicle mixture was disrupted by several freeze-thaw/sonication cycles, followed by extrusion through a polycarbonate filter (100 μm pore size), prior to adding the lipid mixture to the purified SbMATE protein sample, reaching a final lipid concentration of 10 mg/ml and a SbMATE protein concentration between 100 to 300 μg/ml. SbMATE proteo-liposomes were stored at -80 C. Solvent free artificial lipid bilayers were formed on a Port-a-Patch (Nanion) system using commercially available giant unilamellar vesicles (GUVs) preparations and disposable recording glass chips. Upon Giga-ohm seal formation, an asymmetric ionic *cis:trans* gradient (200: 20 mM NaCl pH 7.5) was established across the bilayer and proteoliposomes containing SbMATE were added to the *cis* side.

## ACKNOWLEDGEMENTS

The authors gratefully acknowledgement the contributions of both Quynh Nyugen and J. Michael Green during the production of nanobody binders. APM was supported by an NHMRC Early Career Fellowship (CJ Martin) (GNT1053624). This work was supported by NSF/IOS-1444435 and NIH DK080801.

## REFERENCES

1. Wright, S. H. (2014) Multidrug and Toxin Extrusion Proteins, Wiley Blackwell.

2. Takanashi, K., Shitan, N., and Yazaki, K. (2014) The multidrug and toxic compound extrusion (MATE) family in plants, Plant Biotechnology 31, 417–430.

3. von Uexkll, H. R., and Mutert, E. (1995) Global Extent, Development and Economic-Impact of Acid Soils, Plant and Soil 171, 1–15.

4. Wood S., Sebastian K, and J, S. S. (2000), International Food Policy Research Institute and the World Resources Institute, Washington, DC.

5. Magalhaes, J. V., Liu, J., Guimaraes, C. T., Lana, U. G. P., Alves, V. M. C., Wang, Y. H., Schaffert, R. E., Hoekenga, O. A., Pineros, M. A., Shaff, J. E., Klein, P. E., Carneiro, N. P., Coelho, C. M., Trick, H. N., and Kochian, L. V. (2007) A gene in the multidrug and toxic compound extrusion (MATE) family confers aluminum tolerance in sorghum, Nat. Genet. 39, 1156–1161.

6. Zhou, G., Pereira, J. F., Delhaize, E., Zhou, M., Magalhaes, J. V., and Ryan, P. R. (2014) Enhancing the aluminium tolerance of barley by expressing the citrate transporter genes SbMATE and FRD3, J Exp Bot 65, 2381–2390.

7. Maron, L. G., Guimarães, C. T., Kirst, M., Albert, P. S., Birchler, J. A., Bradbury, P. J., Buckler, E. S., Coluccio, A. E., Danilova, T. V., Kudrna, D., Magalhaes, J. V., Piñeros, M. A., Schatz, M. C., Wing, R. A., and Kochian, L. V. (2013) Aluminum tolerance in maize is associated with higher MATE1 gene copy number, Proceedings of the National Academy of Sciences 110, 5241–5246.

8. Melo, J. O., Lana, U. G. P., Piñeros, M. A., Alves, V. M. C., Guimarães, C. T., Liu, J., Zheng, Y., Zhong, S., Fei, Z., Maron, L. G., Schaffert, R. E., Kochian, L. V., and Magalhaes, J. V. (2013) Incomplete transfer of accessory loci influencing SbMATE expression underlies genetic background effects for aluminum tolerance in sorghum, The Plant Journal 73, 276–288.

9. Yang, X. Y., Yang, J. L., Zhou, Y., Piñeros, M. A., Kochian, L. V., Li, G. X., and Zheng, S. J. (2011) A de novo synthesis citrate transporter, Vigna umbellata multidrug and toxic compound extrusion, implicates in Al-activated citrate efflux in rice bean (Vigna umbellata) root apex, Plant Cell Environ 34, 2138–2148.

10. Yokosho K., Yamaji N., and Ma, J. F. (2010) Isolation and characterisation of two MATE genes in rye, Functional Plant Biology 37, 296–303.

11. Furukawa J., Yamaji N., Wang H., Mitani N., Murata Y., Sato K., Katsuhara M., Takeda K., and Ma, J. F. (2007) An Aluminum-Activated Citrate Transporter in Barley, Plant and Cell Physiology 48 1081–1091.

12. Yokosho K., Yamaji N., and Ma, J. F. (2011) An Al-inducible MATE gene is involved in external detoxification of Al in rice, The Plant Journal 68, 1061–1069.

13. Maron, L., Piñeros, M. A., Guimaraes, C., Magalhaes, J., Pleiman, J., Mao, C., Shaff, J., Belicuas, S. N. J., and Kochian, L. V. (2010) Two functionally distinct members of the MATE (multidrug and toxic compound extrusion) family of transporters potentially underlie two major Al tolerance QTL in maize, The Plant Journal 61, 728-740.

14. Orij, R., Postmus, J., Ter Beek, A., Brul, S., and Smits, G. J. (2009) In vivo measurement of cytosolic and mitochondrial pH using a pH-sensitive GFP derivative in Saccharomyces cerevisiae reveals a relation between intracellular pH and growth, Microbiology 155, 268–278.

15. Zhang, H., Zhao, F.-G., Tang, R.-J., Yu, Y., Song, J., Wang, Y., Li, L., and Luan, S. (2017) Two tonoplast MATE proteins function as turgor-regulating chloride channels in Arabidopsis, Proceedings of the National Academy of Sciences.

16. Doshi, R., Chen, B. R., Vibat, C. R. T., Huang, N., Lee, C.-W., and Chang, G. (2014) In vitro nanobody discovery for integral membrane protein targets, Sci Rep-Uk 4, 6760.

17. Alexandratos, N., and Bruinsma, J. (2012) World agriculture towards 2030/2050: the 2012 revision, In ESA Working paper No. 12–03., FAO, Rome.

18. Carvalho, G., Schaffert, R. E., Malosetti, M., Viana, J. H. M., Menezes, C. B., Silva, L. A., Guimaraes, C. T., Coelho, A. M., Kochian, L. V., van Eeuwijk, F. A., and Magalhaes, J. V. (2016) Back to Acid Soil Fields: The Citrate Transporter SbMATE Is a Major Asset for Sustainable Grain Yield for Sorghum Cultivated on Acid Soils, G3-Genes Genomes Genetics 6, 475–484.

19. Zhou, G. F., Pereira, J. F., Delhaize, E., Zhou, M. X., Magalhaes, J. V., and Ryan, P. R. (2014) Enhancing the aluminium tolerance of barley by expressing the citrate transporter genes SbMATE and FRD3, Journal of Experimental Botany 65, 2381-2390.

20. Omote, H., Hiasa, M., Matsumoto, T., Otsuka, M., and Moriyama, Y. (2006) The MATE proteins as fundamental transporters of metabolic and xenobiotic organic cations, Trends Pharmacol Sci 27, 587–593.

21. Jin Y., Nair A., and van Veen, H. W. (2014) Multidrug transport protein norM from vibrio cholerae simultaneously couples to sodium- and proton-motive force, J Biol Chem 289, 14624–14632.

22. Mazurkiewicz, P., Poelarends, G. J., Driessen, A. J., and Konings, W. N. (2004) Facilitated drug influx by an energy-uncoupled secondary multidrug transporter, J Biol Chem 279, 103–108.

23. Casal, M., Paiva, S., Queiros, O., and Soares-Silva, I. (2008) Transport of carboxylic acids in yeasts, FEMS Microbiol Rev 32, 974–994.

24. Aliverdieva, D. A., and Mamaev, D. V. (2010) Study on the Dicarboxylates Transport into Saccharomyces Cerevisiae Cell Using Its Endogenous Coupled System, Biotech Agr Ind Med, 65–72.

25. Sun, J., Aluvila, S., Kotaria, R., Mayor, J. A., Walters, D. E., and Kaplan, R. S. (2010) Mitochondrial and Plasma Membrane Citrate Transporters: Discovery of Selective Inhibitors and Application to Structure/Function Analysis, Mol Cell Pharmacol 2, 101–110.

26. Velamakanni S., Lau C. H., Gutmann D. A., Venter H., Barrera N. P., Seeger M. A., Woebking B., Matak-Vinkovic D., Balakrishnan L., Yao Y., U, E. C., Shilling, R. A., Robinson, C. V., Thorn, P., and van Veen, H. W. (2009) A multidrug ABC transporter with a taste for salt, PLoS One 4, e6137.

27. He, G. X., Kuroda, T., Mima, T., Morita, Y., Mizushima, T., and Tsuchiya, T. (2004) An H(+)-coupled multidrug efflux pump, PmpM, a member of the MATE family of transporters, from Pseudomonas aeruginosa, J Bacteriol 186, 262–265.

28. Tanaka, Y., Hipolito, C. J., Maturana, A. D., Ito, K., Kuroda, T., Higuchi, T., Katoh, T., Kato, H. E., Hattori, M., Kumazaki, K., Tsukazaki, T., Ishitani, R., Suga, H., and Nureki, O. (2013) Structural basis for the drug extrusion mechanism by a MATE multidrug transporter, Nature 496, 247–251.

29. Lu, M., Radchenko, M., Symersky, J., Nie, R., and Guo, Y. (2013) Structural insights into H+-coupled multidrug extrusion by a MATE transporter, Nat Struct Mol Biol 20, 1310–1317.

30. Pineros, M. A., Cancado, G. M. A., and Kochian, L. V. (2008) Novel properties of the wheat aluminum tolerance organic acid transporter (TaALMT1) revealed by electrophysiological characterization in Xenopus Oocytes: functional and structural implications, Plant Physiol 147, 2131–2146.

